# Gaze biases can reflect task-specific spatial memorization strategies

**DOI:** 10.1101/2024.08.30.610231

**Authors:** Samson Chota, Kabir Arora, J. Leon Kenemans, Surya Gayet, Stefan Van der Stigchel

## Abstract

Previous work has suggested that small directional eye movements not only reveal the focus of external spatial attention towards visible stimuli, but also accompany shifts of internal attention to stimuli in visual working memory (VWM)(van Ede et al., 2019). When the orientations of two bars are memorized and a subsequent retro-cue indicates which orientation needs to be reported, participants’ gaze is systematically biased towards the former location of the cued item (**Figure 1AB**). This finding was interpreted as evidence that the oculomotor system indexes internal attention; that is, attention directed at the location of stimuli that are no longer presented but are maintained in VWM. Importantly, as the location of the bars is presumably not relevant to the memory report, the authors concluded that orientation features in VWM are automatically associated with locations, suggesting that VWM is inherently spatially organized. This conclusion depends on the key assumption that participants indeed memorize and subsequently attend orientation features. Here we re-analyse Experiment 1 by van Ede et al. (2019) and demonstrate that this assumption does not hold. Instead of memorizing orientation features, participants deployed an alternative spatial strategy by memorizing bar endpoints. Although we do not call into question the conclusion that internal attention is inherently spatially organized, our results do imply that directional gaze biases might also reflect attention directed at task-relevant stimulus endpoints, rather than internal attention directed at memorized orientations.

## Analysis

Studies on internal attention via gaze have primarily used rectangular oriented bars (van Ede et al., 2019, 2020; Liu et al., 2022; Draschkow et al., 2022). Bar stimuli are usually rotated around their centre of mass, which means that any orientation maps onto unique endpoints. Therefore, participants could memorize these endpoints instead of the orientation information per se. This strategy could serve as a form of cognitive offloading; akin to keeping a finger pointed at the former stimulus location, maintaining a gaze bias during the delay allows to ‘remember’ a location with minimal mental effort. We tested if participants’ gaze directions depended on the orientation of the stimuli by grouping trials based on the location of the cued item (left versus right) and the orientation of the bars (20 to 70 versus 110 to 160 degree). By doing so, we created four new conditions of interest, in which the nearest endpoint of the memorized bar was either up-right, up-left, down-right, or down-left of fixation (see **Figure 1DEGH**). For each of these endpoint conditions we separately calculated gaze density maps. We then subtracted the average over all maps from each of the four individual maps to highlight their unique contributions. Participants’ gaze showed a significant bias in the direction of the most foveal bar endpoint in all conditions (top-left = -1.08, 95% CI [-1.12, -0.09], top-right = 1.08, 95% CI [0.85, 1.48], down-left = -2.06, 95% CI [-2.73, -2.05], down-right = 2.06, 95% CI [1.76, 2.28], units in radians, **Figure 1C**). Moreover, participants’ gaze was biased away from the left and right horizontal midlines in all conditions (top-left = -1.57, 95% CI [-1.12, -0.09], top-right = 1.57, 95% CI [0.85, 1.48], down-left = -1.57, 95% CI [-2.73, -2.06], down-right = 1.57, 95% CI [1.76, 2.28], **Figure 1C**). Collapsing across orientation conditions separately within the attend left and attend right conditions revealed the originally reported horizontal gaze bias (**Figure 1AB**), whereas collapsing across attention conditions for upwards and downwards oriented bars revealed a strong vertical gaze bias (Rayleigh’s test of uniformity, up: p = 0.004, down: p < 0.0001,**Figure 1FI**). Individual averaged vertical and horizontal gaze biases were significantly correlated, further suggesting that the horizontal gaze biases -at least partly -reflects diagonal biases toward the bar endpoints **(Figure 2E)**. Taken together, these results show that attention, as indexed through gaze biases, was specifically directed to the former location of nearest bar endpoint, rather than the former location of the memorized orientation. We interpret this gaze bias toward the nearest bar endpoint to reflect the strategic substitution of orientation memory by location memory. Evidence for this can already be observed before the presentation of the retro cue: we found vertical gaze biases during stimulus presentation and sustained throughout the entire 2.5-second-long maintenance period until probe presentation (**Figure 2ABD**).

**Figure 1.**
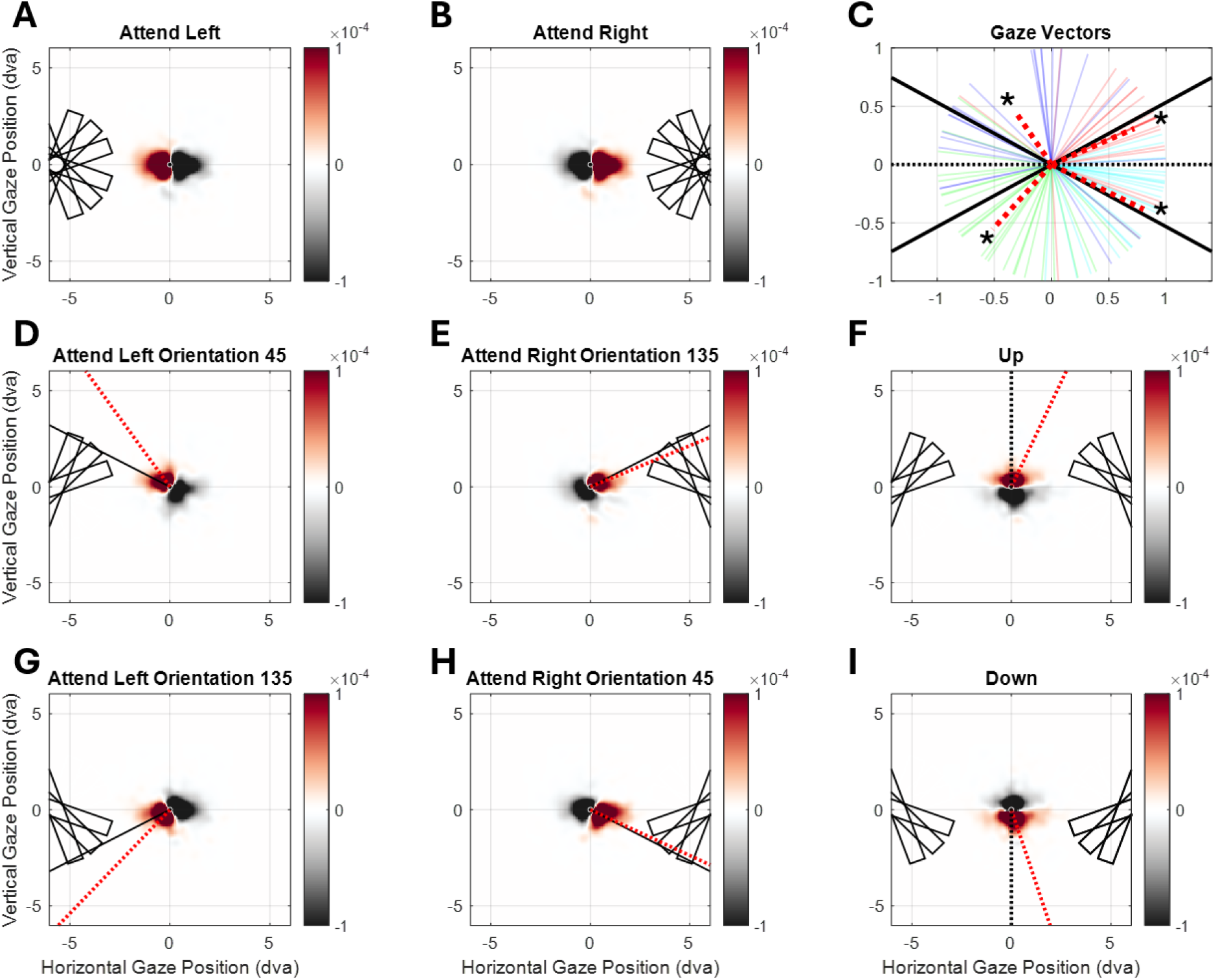
Gaze density maps from Experiment 1 by van Ede et al. (2019) (N = 23, trials included = 20.864, 400 to 1000 ms). **AB**. Original reported effect of cued item location on gaze bias. Calculated by subtracting cued-item-left and cued-item-right gaze density maps. Rectangles indicate used stimulus positions and orientation ranges (min: 20°, mean: 45°, max: 70°; min: 110°, mean: 135°, max: 160°) of bar stimuli. **C**. Normalized Gaze bias vectors per condition (red dotted lines), horizontal vectors (dotted black lines) and average vectors pointing towards most foveal bar endpoints (solid black lines). Gaze bias vector endpoints were calculated from the centre of mass of each condition, ignoring negative values. Circular t-tests revealed that individual gaze bias vector angles (red dotted lines) were significantly different from horizontal vectors (dotted black lines) but not significantly different from endpoint vectors (solid black lines). **FI**. Vertical gaze bias revealed by separating trials based on bar orientations. Red dotted lines depict group average gaze bias vectors. **F**. Both bar endpoints “upwards” (left: 20° to 70° right: 110° to 160°) minus both bars endpoints “downwards” (left: 110° to 160°, right: 20° to 70°). **I**. Both “downwards” minus both “upwards”. **DEGH**. Individual gaze density maps for each attention (left versus right) and bar endpoint direction (upwards versus downwards) separately. Solid black Lines show average vector pointing towards closest 45°/135° bar endpoint (i.e., average optimal gaze location for solving the memory task through memory maintenance of a spatial location). Red dotted lines depict group average gaze bias vectors (calculated from the centre of mass of each condition, ignoring negative values).

**Figure 2.**
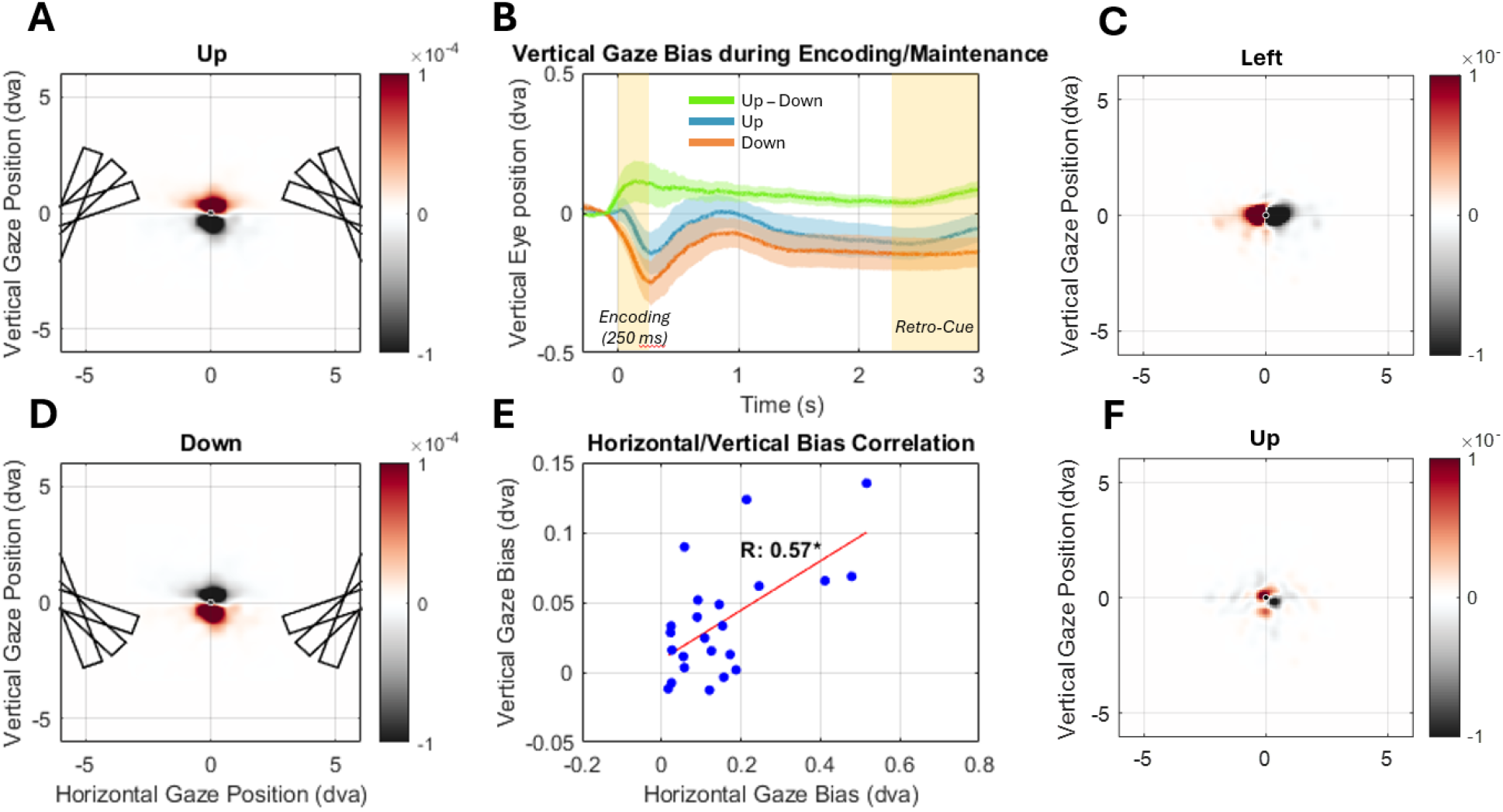
Vertical gaze bias during encoding and maintenance (Experiment 1 by van Ede et al., 2019). **AD**. Gaze density maps during stimulus encoding and maintenance (400 ms to 3000 ms). **B**. Vertical Gaze position during encoding (left yellow shaded area),maintenance and retro-cue (right yellow shaded area) for upwards oriented bars (blue), downwards oriented (orange) bars and the difference (green). Shaded areas show 95% confidence intervals. **C**. Data from Arora et al. (2024). Gaze density maps following retro cue (400 to 1000 ms). Effect of cued item location on gaze bias, calculated by subtracting cued-item-left and cued-item-right gaze density maps. **E**. Inter subject correlation of vertical and horizontal gaze biases (400 ms to 1000 ms). Vertical and horizontal biases were significantly correlated (R = 0.57, p = 0.0043). **F**. Data from Arora et al. (2024). Gaze density maps following retro cue (400 to 1000 ms) reveal no systematic gaze bias depending on Gabor orientation. Calculated by subtracting trials in which both Gabor orientations pointed “downwards” (left: 91° to 180°, right: 1° to 90°) from trials were both orientations pointed “upwards” (left: 1° to 90° right: 91° to 180°).

If participants’ gaze indeed indexed memorized bar endpoints, this further predicts that downwards gaze biases should lead to more clockwise reports in attend left (left-down: negative correlation) and more counterclockwise reports in attend right trials (right-down: positive correlation). Similarly, upwards gaze biases should be associated with more counterclockwise reports in attend left (left-up: positive correlation) and more clockwise reports in attend right trials (right-up: negative correlation). To test this, we performed within-subject trial-by-trial correlations between vertical gaze biases and reported orientations, including only trials with a congruent horizontal gaze bias. Indeed, group-level t-tests on within-subject correlation coefficients indicated positive relationships in left-up (mean rho = 0.01) and right-down (mean rho = 0.05*) conditions and negative relationships in left-down (mean rho = -0.08*) and right-up (mean rho = -0.06*) conditions. This highly specific pattern of correlations between gaze and behaviour further supports the idea that orientation reports were based on memorized endpoint locations.

## Conclusions

Our results have two important implications for the interpretation of gaze biases: First, gaze biases can reflect a stimulus specific spatial strategy that does not involve memorization of orientation per se. Second, the prolonged gaze biases observed before the retro-cue might reflect sustained attention, which has been a matter of recent debate (van Ede et al., 2023; Willet & Mayo, 2023).

Despite these new implications, we do not call into question the main theoretical conclusions of van Ede et al. (2019) that internal attention is inherently spatially organized. Since then, similar gaze biases were observed using stimuli without a clear endpoint (Liu et al., 2024; Ester et al. 2023). In fact, we recently conducted a retro-cue paradigm like that of van Ede et al. (2019) but using oriented Gabors instead of bars (Arora et al., 2024), making a spatial memorization strategy less useful for an orientation recall. Accordingly, we found horizontal gaze biases (reflecting internal attention to the location of the cued item) but no diagonal gaze biases (reflecting a spatial memorization strategy; **Figure 2CF**). Moreover, in their Experiment 4, van Ede et al. (2019) showed that the horizontal gaze bias also emerges when participants need to report the color of a bar that was cued via its orientation (0° vs 90°). In this case, maintaining attention (or gaze) toward the nearest bar endpoint is not a useful strategy for the task of reporting the color. In sum, there is ample support for the original conclusions of van Ede et al. (2019) that visual working memory is inherently spatially organized.

To conclude, we call for caution in interpreting gaze biases, akin to those reported in the article of van Ede et al. (2019), as unequivocally reflecting shifts of internal attention. Particular stimuli and task designs may allow for alternative (e.g., spatial) memorization strategies. Furthermore, and theoretically of greater interest, our results show that observers can approach even the simplest working memory recall tasks in a variety of ways. When set-up and stimuli allow for, observers may deploy unexpected but resource-efficient strategies, highlighting the richness of visuospatial working memory use.

## Author contributions

S.C. and K.A. analysed the data. S.C., K.A., S.G., J.L.K and S.v.d.S. wrote and edited the paper. We want to thank Freek van der Ede for his help with data analysis and advise for the manuscript.

## Data availability

The authors of the original paper have made their data available through the Dryad Digital Repository at: https://doi.org/10.5061/dryad.m99r286.

## Code accessibility

Code is available from the authors on reasonable request.

